# Revealing the critical regulators of cell identity in the mouse cell atlas

**DOI:** 10.1101/412742

**Authors:** Shengbao Suo, Qian Zhu, Assieh Saadatpour, Lijiang Fei, Guoji Guo, Guo-Cheng Yuan

**Affiliations:** Department of Biostatistics and Computational Biology, Dana-Farber Cancer Institute and Harvard T. H. Chan School of Public Health, Boston, MA 02215, USA; Center for Stem Cell and Regenerative Medicine, Zhejiang University School of Medicine, Hangzhou, 310058, China

## Abstract

Recent progress in single-cell technologies has enabled the identification of all major cell types in mouse. However, for most cell types, the regulatory mechanism underlying their identity remains poorly understood. By computational analysis of the recently published mouse cell atlas data, we have identified 202 gene regulatory networks whose activities are highly variable across different cell types, and more importantly, predicted a small set of essential regulators for each of over 800 cell types in mouse for the first time. Systematic validation by automated literature- and data-mining provides strong additional support for our predictions. Thus, these predictions serve as a valuable resource that would be useful for the broad biological community. Finally, we have built a user-friendly, interactive, web-portal to enable users to navigate this mouse cell network atlas.

## Introduction

A multi-cellular organism contains diverse cell types, each has its own functions and morphology. A fundamental goal in biology is to characterize the entire cell-type atlas in human and model organisms. With the rapid development of single-cell technologies, great strides have been made in the past few years (Blainey and Quake, 2014; Sandberg, 2014; Stegle et al., 2015; Svensson et al., 2018; Yuan et al., 2017). Multiple groups have made tremendous progresses in mapping cell atlases in complex organs (such as mouse brain and immune system) (Rosenberg et al., 2018; Stubbington et al., 2017), early embryos (such as in *C elegans* and zebrafish) (Cao et al., 2017; Wagner et al., 2018), or even entire adult animals (such as *Schmidtea mediterranea* and mouse) (Fincher et al., 2018; Han et al., 2018; Plass et al., 2018). International collaborative efforts are underway to map out the cell atlas in human (Regev et al., 2017).

How do cells maintain their identity? While it is clear the maintenance of cell identity involves the coordinated action of many regulators, and transcription factors (TFs) have been long recognized to play a central role. In several cases, the activity of a small number of key TFs, also known as the master regulators, are essential for cell identity maintenance: depletion of these regulators cause significant alteration of cell identity, while forced expression of these regulators can effectively reprogram cells to a different cell-type (Han et al., 2012; Ieda et al., 2010; Riddell et al., 2014; Takahashi and Yamanaka, 2006). However, for most cell-types, the underlying gene regulatory circuitry is incompletely understood. With the increasing diversity of gene expression programs being identified through single-cell analysis, an urgent need is to understand how these programs are established during development, and to identify the key regulators responsible for such processes.

Systematic approaches for mapping gene regulatory networks (GRNs) have been well established. The most direct approach is through genome-wide occupancy analysis, using experimental assays such as ChIPseq, chromatin accessibility, or long-range chromatin interaction assays (Consortium, 2012). However, this approach is not scalable to a large number of cell types, and its application is often limited by the number of cells that can be obtained *in vivo*. An alternative, more generalizable approach is to computationally reconstruct GRNs based on single-cell gene expression data (Friedman, 2004; Gerstein et al., 2012; Karlebach and Shamir, 2008), followed by more focused experimental validations. In this study, we took this latter approach to build the first comprehensive cell network atlas in mouse.

To this end, we took advantage of the recently mapped Mouse Cell Atlas (MCA) derived from comprehensive single-cell transcriptomic analysis (Han et al., 2018), and combined with a computational algorithm to constructing GRNs from single-cell transcriptomic data. Our analysis indicates that most cell-types have distinct regulatory network structure and identifies regulators that are critical for cell identity. In addition, we provide an interactive web-based portal for exploring the mouse cell network atlas.

## Results

### Reconstructing gene regulatory networks using the mouse cell atlas

A number of computational methods have been developed to predict gene regulatory networks from single-cell gene expression data (Friedman, 2004; Gerstein et al., 2012; Karlebach and Shamir, 2008). Recently, the SCENIC method has emerged as a powerful approach for constructing GRNs from single-cell RNAseq data(Aibar et al., 2017; Davie et al., 2018). In brief, It links *cis*-regulatory sequence information together with single-cell mRNA-seq data to identify a list of regulons (each representing a TF along with a set of co-expressed and motif significantly enriched target genes), and the regulon activity scores (RAS) for each cell (see Methods). However, previously such analysis has only been applied to dissect the GRNs in a small number of cell types (Dong et al., 2018; Wuidart et al., 2018). One of the challenges for applying SCENIC to analyze large datasets such as the mouse cell atlas is the scalability. In addition, dropout events are extensive in single-cell mRNA-seq data, and their presence may lead to significant error in computational inference.

To overcome these challenges, we modified SCENIC to enhance robustness and scalability. In particular, we divided the entire cell population into small groups, each containing 20 cells, by random selection from the same tissue and cell-type (Figure 1A) with the assumption that the identified cell types should be biological replicates and that any slight difference within cell type might be technical noise. This step significantly increased the scalability of SCENIC. To evaluate its performance, we compared the results for several representative tissues, each containing multiple cell types. We found that our modified approach, which we refer to as Avg20, separates different cell types more effectively compared to the original implementation (Figure S1). To further evaluate the robustness of our Avg20 approach, we repeated the procedure three times, each creating a different random sample. We found that the results are highly consistent (Figure S1). Taking the bladder tissue for example, the TFs of regulons among different replicates are significantly overlapped (*p*<1e-22, Fisher’s exact test) and the correlations for different replicates larger than 0.84 (see Methods).

**Figure 1.**
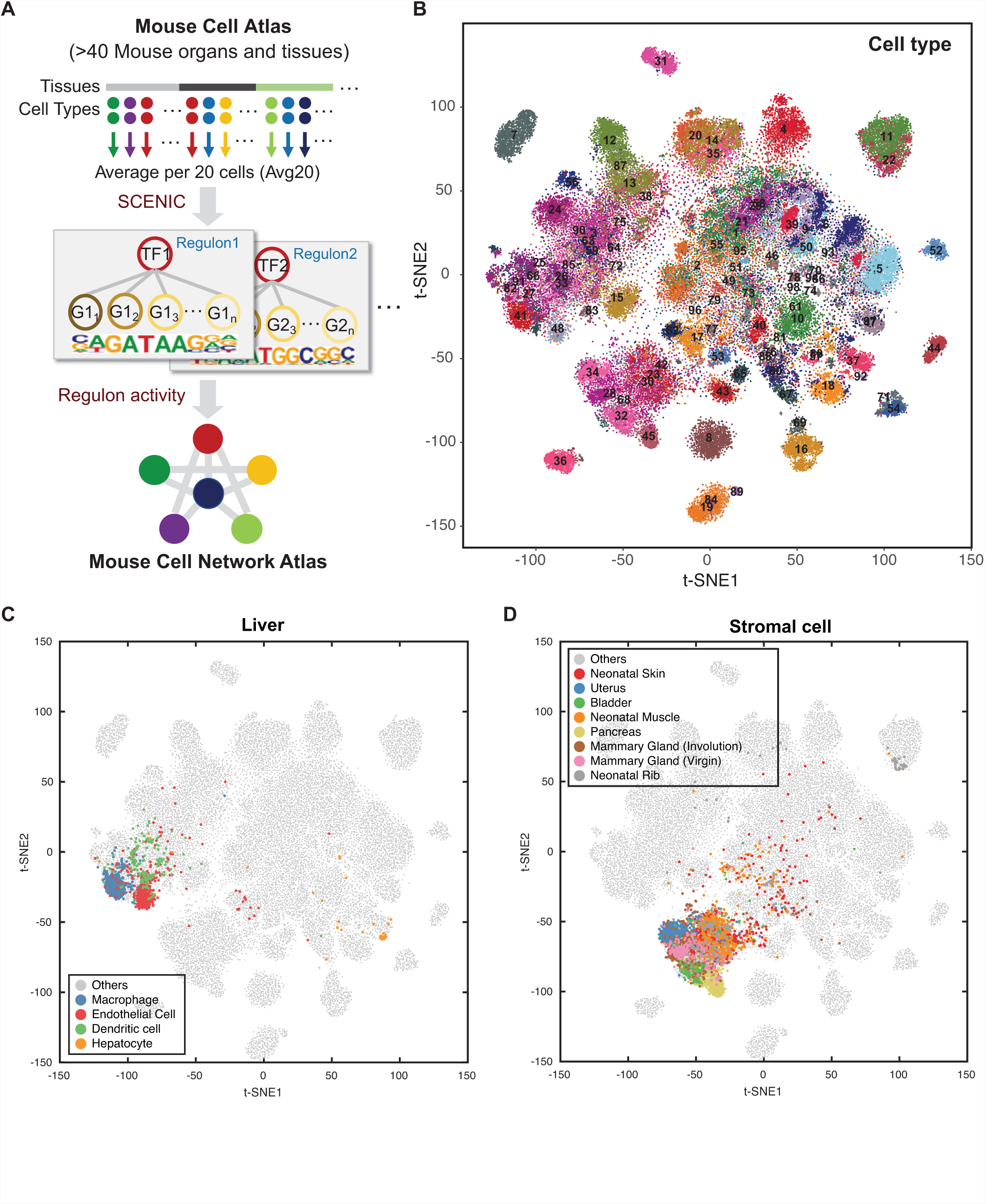
Mapping mouse cell network atlas with regulon activity. (A) Schematic overview of our approach. Expression values are first averaged per 20 cells (Avg20) within each cell type and tissue. Regulons are then identified by SCENIC based on averaged expression values. Regulon activity scores are calculated for each individual cell. Cell type relationship is quantified based on the similarity of regulon activities. (B-D) t-SNE map for all sampled single cells (∼61k) based RAS, each cell is color-coded based on cell-type assignment. (B) all sampled cells (61k) are highlighted. (C) Liver cells are highlighted. (D) Stromal cells are highlighted.

Next, we considered a representative subset of MCA data (Han et al., 2018), containing 61,637 cells sampled from 43 tissues. Previous analysis has identified 98 main cell-types (Han et al., 2018). By applying the Avg20 approach described above, we identified 202 significant regulons containing 8,461 genes (Table S1). Here, we also repeated the Avg20 approach three times to further test the robustness based on this big dataset and got very high agreement (Figure S2A, *p*<1e-41, correlation between replicates>0.94). For comparison, we repeated this analysis but using a different group size, corresponding to 50 cells (Avg50) and 100 cells (Avg100), respectively. While the overall performance is about the same, we observed a slight reduction of efficiency for separating cell types between different random samples (*p*<1e-37, Fisher’s exact test, Figure S2B). Therefore, we settled with using the Avg20 approach for the rest of this study, unless otherwise stated.

The size of each regulon varies from 10 to 2,502 genes, with a median size of 73 genes. Strikingly, different cell-types are well separated using the RAS-based distance (Figure 1B, Figure S3A, Table S2). Even within a single tissue, different cell-types can be separated based on the regulon activities (Figure 1C, Figure S3B). For example, each of four main cell-types identified in liver occupies a distinct territory in the tSNE plot (Figure 1C).

One question of interest is whether cells of the same type may have different regulatory circuitries across tissues. Such differences would be relevant for investigating cell-environment interactions (Figure 1D, Figure S3C). To address this question, we focused on stromal cells, which can be found in a wide variety of tissues, providing support, structural and anchoring functions. The behavior of stromal cells is well known to be highly plastic, a necessary property for supporting a diverse range of tissue development (Lee et al., 2006). Indeed, we found that the stromal cells from different tissues tend to have distinct regulon activities (Figure 1D). Specifically, while stromal cells are clustered together from the global view of t-SNE map, closer examination suggests the subpopulations from different tissues, such as uterus, mammary gland, bladder and pancreas, are well separated. Similar refined structure can be found in various other cell types, such as T cells and epithelial cells (Figure S3C). Taken together, our cell type-based strategy for single cell analysis provides a new avenue to investigate the potential regulatory mechanism in inter- and intra-cell type variations. The results also indicates that GRN differences are primarily driven by cell-type differences, but further modulated by tissue environment differences.

### Comparative analysis identified essential regulators for the maintenance of cell identity

Our comprehensive network analysis provides an opportunity to systematically identify critical regulators for cell identity. For each regulon, we evaluated its activities associated with each of the 98 major cell types (Table S3), and defined a regulon specificity score (RSS) based on Jensen-Shannon divergence (Cabili et al., 2011) (Table S4, see Methods). We then selected the regulons with highest RSS values and further examined their functional properties. To test whether this approach is effective, we started with the erythroblast because its core gene regulatory network has been well characterized (Cantor and Orkin, 2002). Our network analysis identified *Lmo2*, *Gata1* and *Tal1* (also known as SCL), as the most specific regulons associated with erythroblast (Figure 2A). tSNE plot provides additional support that the activities of these regulons are highly specific to erythroblast (Figure 2B-C). Of note, all three factors are well-known master regulators for erythrocytes (Welch et al., 2004; Wilson et al., 2010; Wu et al., 2014). Another well-characterized cell-type is the B cell. Our network analysis identified *Ebf1* and *Bcl11a* as the most specific regulons (Figure 2E-G). Both factors are well known to be essential regulators for maintaining B cell identity (Nechanitzky et al., 2013), (Liu et al., 2003).

**Figure 2.**
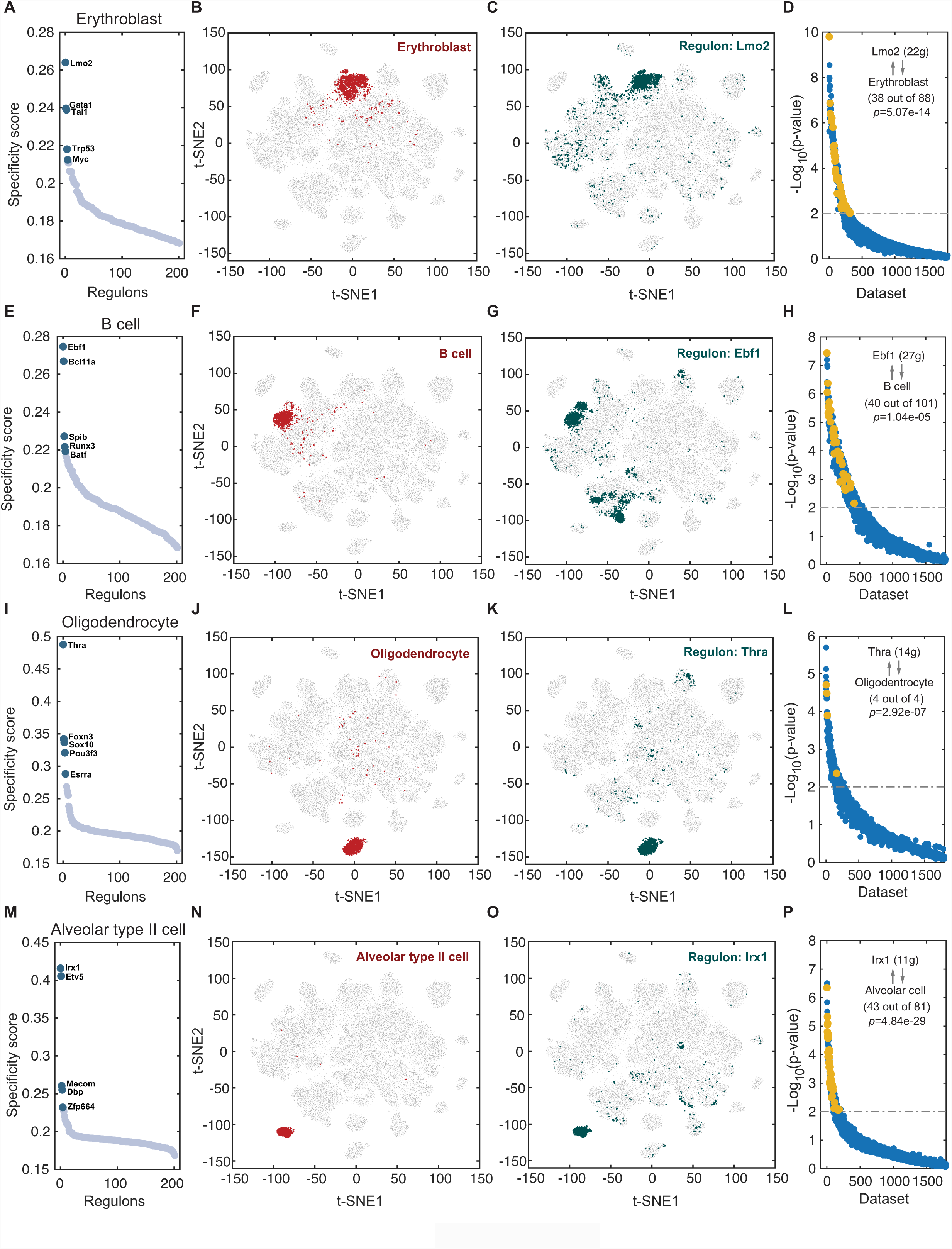
Cell-type specific regulon activity analysis. (A-D) Erythroblast. (A) Rank for regulons in erythroblast cell based on RSS. (B) Erythroblast cells are highlighted in the t-SNE map (red dots). (C) Binarized RAS (use *z*-score value of 2.5 as cutoff) for top regulon *Lmo2* on t-SNE map (dark green dots). (D) SEEK co-expression result for target genes of top regulon *Lmo2* in different GEO datasets. The x-axis represents different datasets, and the y-axis represents the significance of target genes in each dataset. Erythroblast related datasets with correlation *p* value<0.01 are highlighted by yellow dots. (E-H) Same as (A-D) but for B-cells. (I-L) Same as (A-D) but for oligodendrocytes. (M-P) Same as (A-D) but for Alveolar type II cells.

The success of our approach in recapitulating critical regulators for well-characterized cell-types motivated us to repeat the analysis for all other 96 cell-types. For each cell-type, we identified a small number of regulons with exceptionally high specificity scores (Figure 2, Figure S4). To systematically evaluate the accuracy of these predictions, we used two complementary approaches: SEEK (Zhu et al., 2015) and CoCiter (Qiao et al., 2013), based on mining the pubic datasets and literatures, respectively. First, SEEK analysis was done to select datasets in which the TF and its target genes within a regulon are co-expressed. We queried the titles for public mouse datasets for enrichment of cell-type specific terms, with the assumption that functionally related genes tend to be co-expressed in the corresponding cell-types. Second, CoCiter analysis was done to identify enriched co-occurrence of a gene and cell-type term pair in publication abstracts, with the assumption that functionally related genes and terms should frequently appear together in the literature.

To test if the above two data-mining approaches are useful validation strategies, we applied each approach to test the regulators identified for erythroblast and B cells. For erythroblast, we applied SEEK analysis to search for GEO datasets in which the genes in regulon *Lmo2* are significantly co-expressed. Among the more than 2,000 datasets examined by SEEK, the erythroblast related datasets are highly ranked (Figure 2D, Table S5, Fisher’s exact test, *p*=5.07e-14, see Methods). Similarly, we applied CoCiter analysis to search for publications where genes in regulon *Lmo2* co-occur with the term ‘erythroblast/erythrocyte’. Again, the result is highly specific (Co-citation Impact, CI=8.41, permutation *p*<0.001, see Methods). Similarly, for B cells, the top ranked datasets in which *Ebf1* and its target genes are co-expressed tend to be associated with B cells (Figure 2H, *p*= 1.04e-05), and these genes tend to co-occur in studies related to B cells (CI=9.38, permutation *p*<0.005).

The above analysis indicates that both SEEK and CoCiter analyses are useful for validating predicted essential regulators of cell identity. Therefore, we applied both approaches to evaluate the relevance of predicted essential regulators for other cell-types, most of which are incompletely characterized. For example, oligodendrocytes are a type of neuroglia whose main functions are to provide support and insulation to axons in the central nervous system. While various factors have been implicated to play a role in oligodendrocyte development (Zuchero and Barres, 2013), the most important regulators remain unknown. Our network analysis found that regulon *Thra* shows the highest specificity score (Figure 2I-K), suggesting this may be one of the most essential regulators for oligodendrocyte identity. As additional support, SEEK analysis indicated that the genes in regulon *Thra* are significantly co-expressed in oligodendrocyte related datasets (Figure 2L, *p*= 2.92e-07). For the potential target genes of *Thra*, CoCiter analysis shows that there are 204 papers (CI=7.68, permutation *p*<0.005) mentioned both the genes of regulon *Thra* and ‘oligodendrocyte’ in their abstracts.

Approximately 15 % of lung cells belong to Alveolar type II (AT2), which has the important functions of synthesizing and secreting surfactant (Mason, 2006). Our GRN analysis indicates that regulon *Irx1* shows the highest specificity score in these cells (Figure 2M-O), indicating this TF may be an essential regulator for AT2 cell identity. As additional support, SEEK analysis shows the genes in regulon *Irx1* significantly co-express in alveolar related datasets (Figure 2P, *p*=4.84e-29). Also, literature mining finds that the genes in this regulon are associated with ‘alveolar’ (CI=8.02, permutation *p-*value<0.001). Taken together, these results strongly indicate that our predicted cell-type specific regulators are functionally relevant.

### Regulons are organized into combinatorial modules

Transcription factors often work in combination to coordinate gene expression levels. To systematically characterize the combinatorial patterns, we compared the atlas-wide similarity of RAS scores of every regulon pair based on the Connection Specificity Index (CSI) (Fuxman Bass et al., 2013) (see Methods). Strikingly, these 202 regulons are organized into 8 major modules (Figure 3A, Figure S5A, Table S6). For each module, we identified several representative regulators and cell types through their average activity scores (Figure S5B). When mapping the average activity score of each module onto tSNE map, we found that each module occupies distinct region and all highlighted regions show complimentary patterns (Figure S5C). Module M1 contains regulators *Gata1, Tal1*, and *Lmo2*, which are essential regulators for the erythroblast. M2 contains regulators that are associated with epithelial proliferation and endothelial apoptosis, such as *Jun* and *Fos* (Okada et al., 2004; Wang et al., 1999). M3 contains a mixture of stroma-specific factors, such as *Twist2* and *Nfix* (Galvan et al., 2015; Grabowska et al., 2014), as well as regulators for bone formation, such as *Creb* family members (Murakami et al., 2009). Several M4 regulators, such as *Ppard* (Shi et al., 2017) and *Klf3* (Kaga et al., 2017), are associated with epithelial cells. Module M5 contains regulators that are specifically activated in testicular cells, such as *Sox5* (Kiselak et al., 2010) and *Ovol2* (Chizaki et al., 2012). Regulons in M6 are highly associated with the nervous system, such as oligodendrocytes and astrocytes. The activity of M7 including *Mafb*, *Irf2* and *Nfkb1* is specifically high in different immune cell types (Valledor et al., 1998), such as macrophages, microglia, dendritic cells, B cells and T cells. Module M8 is also related to immune cells but it is very specifically enriched in one subtype of T cells which are derived from the thymus tissue (Cluster 8, Figure 1B, Figure S5C). A closer examination of this cell type indicates that the regulon *Rorc* is specifically activated in this subtype. It is known that RORγt (encoded by *Rorc*) is essential for T cell maturation in the thymus and could suppress conventional effector responses such as proliferation and cytokine production (Sun et al., 2000; Yui and Rothenberg, 2014), which indicates this subtype of T cells might be in a ‘naïve’ state. In contrast, the T cell subtypes that are associated with M7 have distinct regulon activities and reside in other tissues. For example, T cells from Clusters 3 and 15 are mainly from mammary gland tissues. In these subtypes, the regulon *Batf* (contained in M7) has the highest specificity score. Since *Batf* has been identified to play a fundamental role in regulating the differentiation of effector of CD8+ T cells (Kurachi et al., 2014), these two T cell subtypes are likely to be in an ‘activated’ state.

**Figure 3.**
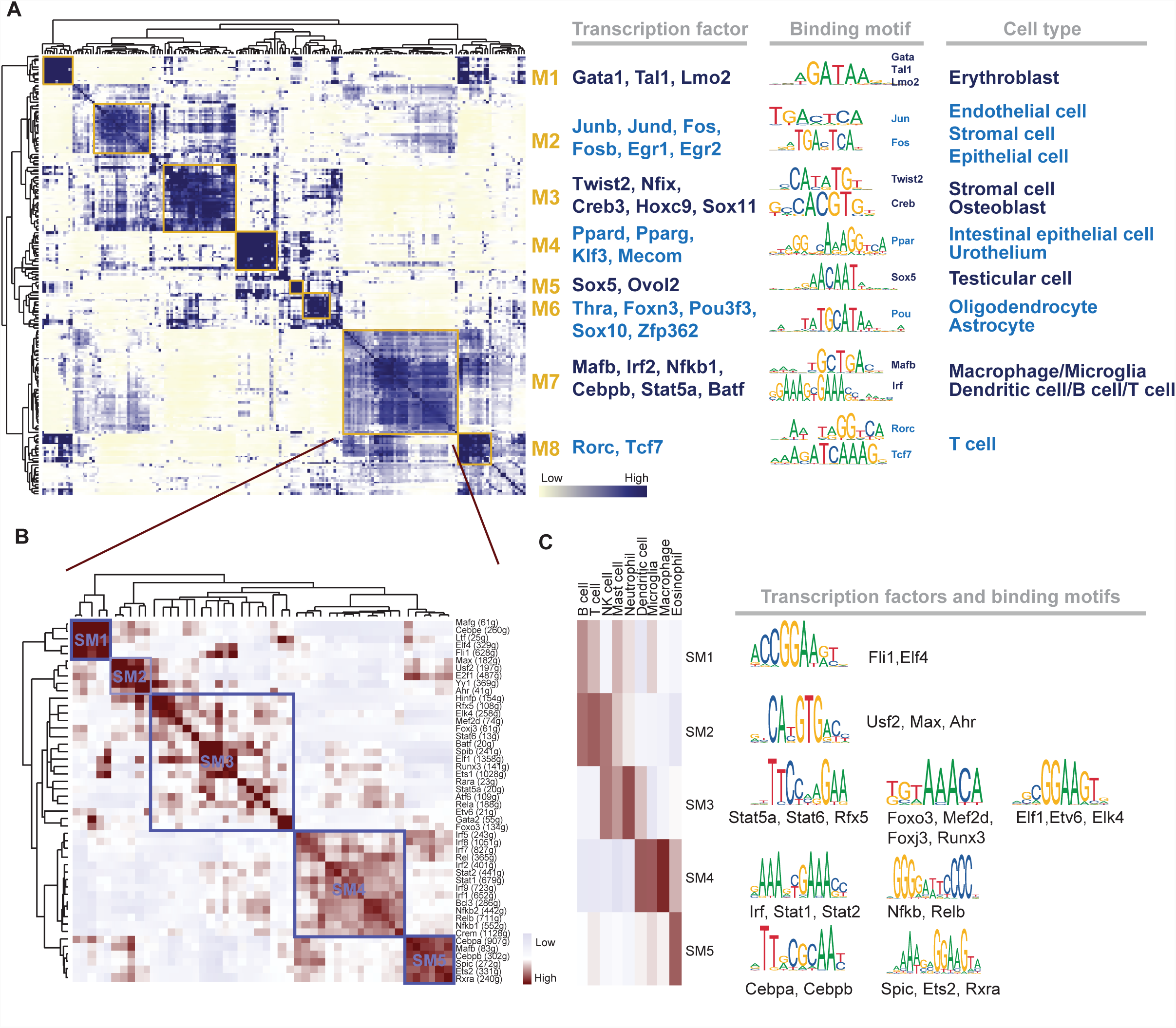
Identification of regulon modules. (A) Identified regulon modules based on regulon CSI matrix, along with representative TFs, corresponding binding motifs and associated cell types. (B) Zoomed-in view of module M7 identifies submodule structures. (C) Different sub-modules in M7 are associated with distinct immune cell types and regulon activities.

We next focused on the largest module M7, which contains 48 regulons. This is perhaps not surprising, considering the complexity of the mammalian immune system. Consistent with this observation, the target genes in these regulons are enriched for immune related functions, such as ‘defense response’, and ‘immune system response’. A closer look reveals that M7 can be further divided in 5 smaller sub-modules (SM1-5) (Figure 3B). Interestingly, each smaller module is specifically associated with distinct immune cell types (Figure 3C). For example, SM3 is associated with neutrophils, and the involved regulons including *Stat5a* (Fievez et al., 2007), *Foxo3* (Osswald et al., 2018) and *Elf1*(Bjerregaard et al., 2003) are associated with neutrophil regulation. SM4 is mainly related to macrophages; correspondingly, many well-known macrophage related regulators are predicted by our network analysis, such as *Irf* family genes and *Nfkb* genes (Valledor et al., 1998). Regulons in SM5 are highly activated in eosinophil cells, and it has been proved that C/EBP genes are required for eosinophil lineage commitment and maturation (Nerlov et al., 1998). These analyses indicate a hierarchical organization of regulatory modules that work together in fine-tuning cellular states.

### Mapping the mouse cell atlas network using cell-type specific regulatory activity

The full MCA dataset contains over 800 cell types (Table S7). At this resolution, many cell types share similar gene expression patterns and their biological functions are likely to be less distinct. Our complete network analysis estimated the RAS (Table S8) and RSS (Table S9) for all these cell types, while related cell-types share similar overall network structure. Such relationship is represented by a highly modularized graph, where each edge connects a pair of related cell types whose overall regulon activities is similar (Spearman correlation > 0.8, Figure 4A). The network global structure is similar to that of the 98 major cell types (Figure S6) This graph can be further divided into 21 groups (G1-G21) by using the Markov Clustering Algorithm (MCL) (See Methods), with functional related cell types are clustered together. 9 of these groups (G1-G9) contain more than 10 related cell types (Figure 4A-B). For example, G1 contains a number of blood related cell-types, whereas cell-types in G2 perform various supporting functions. The Sankey plot (Figure 4B) summarizes the relationship between cell types and their top associated regulon modules. For example, M1, M7 and M8 are highly enriched in G1, whereas modules M2 and M3 are highly enriched in G2.

**Figure 4.**
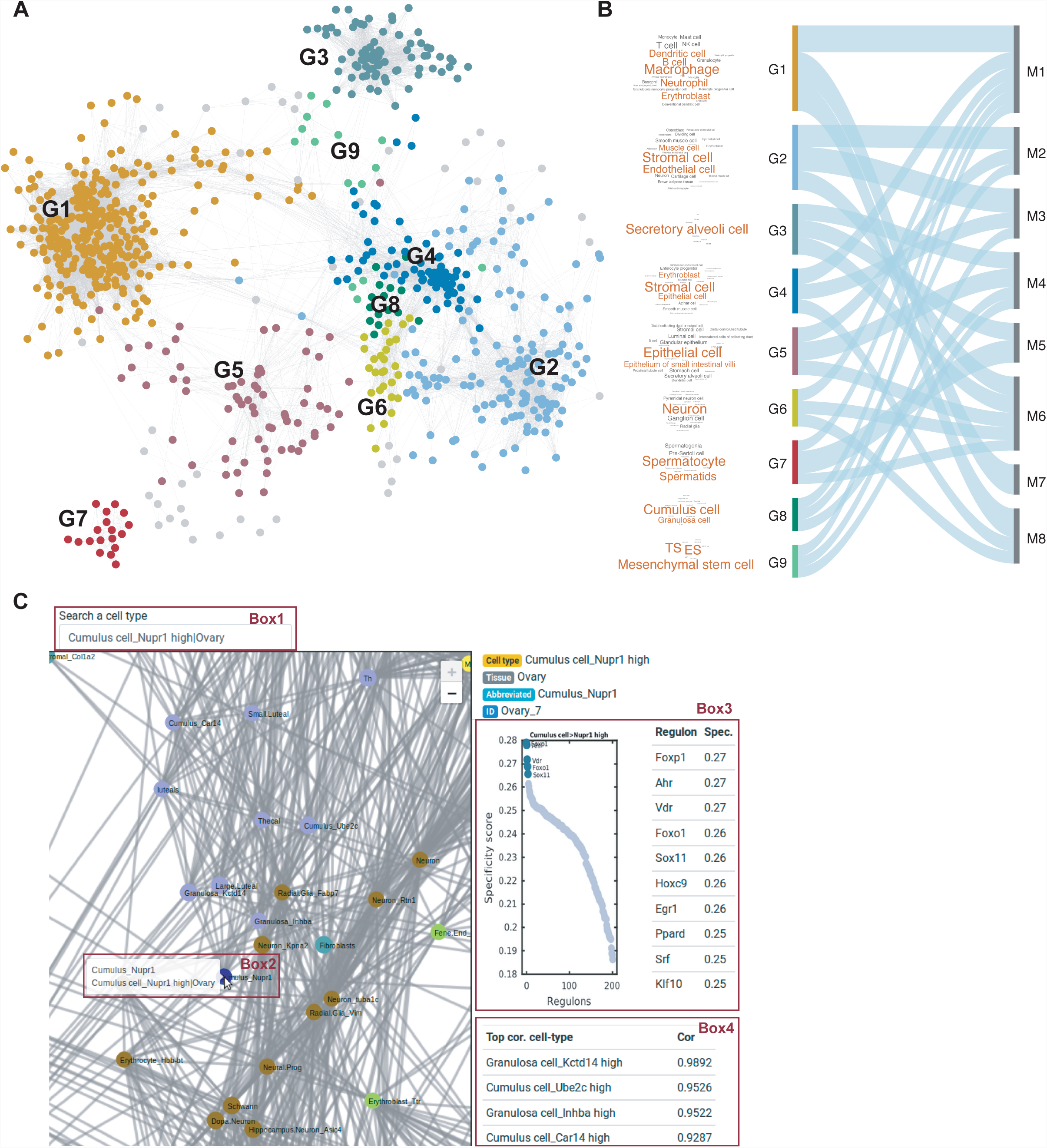
A summary view of the mouse cell network atlas. (A) Relatedness network for the 818 cell types based on similarity of regulon activities. Each group represents highly related cell types. (B) Sankey plot shows relationship between cell-type groups G1-9 and regulon modules M1-8, the thematic cell type composition within each cluster is indicated by the corresponding wordcloud plot. (C) A representative screenshot of the web-portal obtained by querying ‘cumulus cells’.

### A web-based resource for interrogating mouse cell network atlas

We created a web-based portal to enable the users to easily navigate this mouse cell network atlas (http://regulon.rc.fas.harvard.edu/). The web interface allows users to easily interact with the cell type network and the regulon network. We provide two versions of the cell type network, corresponding to 98 major cell types (which are analyzed here) and 818 cell types (from the whole mouse cell atlas), respectively. The regulon network represents the relationship between the 202 regulons; each edge connects a pair of regulons whose cell-type specific activities are highly correlated. Users can query a cell type of interest to get information about its associated regulons as well as neighboring cell types in terms of the network structure. For example, cumulus cells are a specific cell type in the ovary whose function is not well characterized. To find information about this cell-type, a user could simply enter “cumulus cell” in the search box to search for related cell types (Figure 4C, Box 1). This will lead to a menu containing related cell types identified by the MCA, one of these cell-types is encoded as “Cumulus cell_Nupr1 high” (Figure 4C, Box2), as *Nupr1* is a distinct marker for this cell type. Selecting this cell type would generate two main outputs. The first output is a list of regulons ranked by the degree of RSS. The most specific regulons are *Foxp1*, *Arh* and *Vdr,* although a number of additional regulons, such as *Foxo1,* have similar specificity (Figure 4C, Box 3). The second output is a list of cell types with similar regulatory networks. Not surprisingly, its neighboring cell-types include other cumulus cell subtypes identified by MCA. Of interest, granulosa cells, which also called cumulus granulosa cells depending on location within the ovarian follicle, are also identified as its neighbors (Figure 4C, Box 4). Thus, biomedical investigators can identify putative regulons that are likely important during mouse ovary and oocyte development. The web-portal also provides a zoom function, which enables users to interactively explore the regulon and cell-type network structures at any desired resolution. In addition, the raw data are also downloadable from the web-portal to support further investigation.

## Discussions

Our knowledge of the cellular heterogeneity has exploded in the past few years. In comparison, for most of the cell-types identified so far, we lack the mechanistic understanding how their characteristic gene expression programs are established and maintained. Neither do we understand the developmental and functional relationship between different cell types. Such information is not just of fundamental biological interest, but also can guide developing novel cell reprogramming strategies and have clinical implications. Building upon the recently mapped mouse cell atlas, we have comprehensively constructed the GRNs for all major cell types in mouse through computational analysis. An important consequence is the predictions of critical regulators for each individual cell type. While most predictions remain hypotheses, they have provided a guide for future experimental investigation. As such, we have created a valuable resource for the broad biology community.

## Methods

### Data sets

The MCA data set and the corresponding cell type annotation were downloaded from https://figshare.com/s/865e694ad06d5857db4b. It contains two parts: (1) the subsampled ∼ 61K single cells date set, which includes 98 major cell types and covers 43 tissues and organs; (2) the whole MCA data set, which contains over 250K single cells after removing the low-quality cells mapped to 818 cell types.

### Inference of regulons and their activity

SCENIC (Aibar et al., 2017) was used to infer the single cell regulatory network (regulon) with modification as described below. Briefly, the regulon inference mainly contains the following three steps: (1) identify co-expression modules between TF and the potential target genes; (2) for each co-expression module, infer direct target genes based on those potential targets for which the motif of the corresponding TF is significantly enriched. Each regulon is then defined as a TF and its direct target genes; (3) the RAS in each single cell is calculated through the area under the recovery curve (see ref. 19 for details).

The original implementation of SCENIC is not scalable to large datasets and its results can be significantly affected by sequencing depth. To improve the scalability or robustness, we modified it by pooling data from every 20 cells randomly selected within each cell type and tissue. Then we applied SCENIC to the pooled data. This simple modification (referred to as Avg20) effectively increases the data quality as well as reduces the computational burden.

We compared the performance of our modified approach with the original version of SCENIC by analyzing the MCA data from three representative tissues: bladder, kidney and bone marrow. For each tissue, the Avg20 approach was repeated three times to estimate the variability due to random sampling. To evaluate the performance of each method, we calculated the Silhouette value, which is a commonly used quantitative metric for clustering consistency. Specifically, the silhouette value *S*_*i*_ for the *i*th cell is defined as *S*_*i*_ = (*b*_*i*_-*a*_*i*_)/max(*b*_*i*_,*a*_*i*_), where *a*_*i*_ is the average distance between the *i*th single cell and other single cells from the same cell type, and *b*_*i*_ is the minimum average distance from the *i*th single cell to cells in a different cell type. A high silhouette value indicates that the cells are well matched to its own cell type, which further indicates that this regulon activity matrix is much better to interpret the cell identity. The results are shown in Figure S1. The consistency between three replicates was evaluated by the following. First, the overlap among the TFs of regulons was evaluated by Fisher’s exact test. Second, for each Avg20 replicate, we also calculated the pairwise distance of single cells in each tissue based on their RAS and then calculated Pearson correlation coefficient (PCC) to evaluate the agreement of different Avg20 replicates. The same strategy was used to evaluate the performance of alternative pooling options: Avg50 and Avg100.

### Quantifying cell-type specific by using regulon specificity score

To quantify the cell-type specificity of a regulon, we used a similar strategy that was previously used for gene expression data analysis (Cabili et al., 2011) to define an entropy-based specificity score, as follows. First, we use a vector 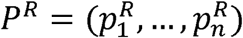 to represent the distribution of RAS in the cell population (*n* is the total number of cells). Here, the RAS are normalized so that 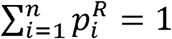 Then, we use a vector 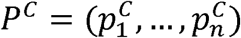 to indicate whether a cell belongs to a specific cell-type 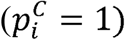 or not 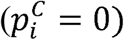 This vector is also normalized. Next, we define the cell-type specificity score by using the Jensen-Shannon Divergence (JSD), which is a commonly used metric for quantifying the difference between two probability distributions, defined as

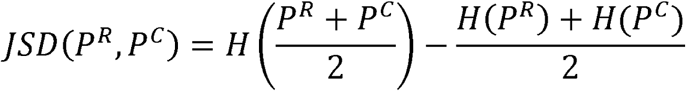

where H(P)=-Σ *p*_*i*_ log (*p* _*i*_)r logcr J represents the Shannon entropy of a probability distribution P. The range of JSD values is between 0 and 1, where 0 means identical distribution and 1 means extreme difference. Finally, the regulon specificity score (RSS) is defined by converting JSD to a similarity score:

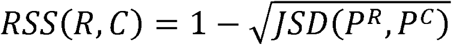

For each cell type, the essential regulators are predicted as those associated with the highest cell-type specific scores.

### Validate the potential function of regulons

We apply the following two methods to validate whether the genes in regulon are related to the function of a certain cell type: (1) SEEK analysis (Zhu et al., 2015), and (2) CoCiter analysis (Qiao et al., 2013). First, SEEK (http://seek.princeton.edu/modSeek/mouse/) is a tool that provides gene co-expression search function for over ∼2000 mouse datasets from the Gene Expression Omnibus (GEO). We used the mouse version of SEEK to evaluate whether the genes in a regulon are co-expressed, and if so whether the datasets supporting the co-expression are associated with an interested cell type. If genes are significantly co-expressed in many datasets related to a certain cell type, it could be inferred that the function of this regulon is highly related to this cell type. Taking the erythroblast for example, we input gene list of regulon *Lmo2* to SEEK web server and search for erythroblast related keywords (such as ‘erythroblast’, ‘hematopoietic’, *etc*.) from the complete dataset-list ranked by query-coexpression score. Then we choose p<0.01 to select significant datasets and finally use Fisher’s exact test to evaluate whether the selected datasets are significantly enriched in the top ranks. Second, the CoCiter (Qiao et al., 2013) is a text mining approach against the up-to-date Medical Literature Analysis and Retrieval System Online (MEDLINE) literature database to evaluate the co-citation impact (CI, log-transformed paper count) between a gene list and a term. To assess significance of co-citation, a Monte Carlo approach is used to evaluate random expectations by randomly selecting 1000 gene sets with the same size as input gene list and then a permutation *p* value is calculated as the number of times that *CI*_*random*_>*CI*_*true*_ divided by 1000. Here we used the function “gene-term” in CoCiter (use default parameters but set organism as mouse, www.picb.ac.cn/hanlab/cociter) to check whether the genes in regulon are significantly co-cited with a certain cell type in literatures.

### Regulon module analysis

Regulon modules were identified based on the Connection Specificity Index (CSI) (Fuxman Bass et al., 2013), which is a context-dependent measure for identifying specific associating partners. The evaluation of CSI involves two steps. First, the Pearson correlation coefficient (PCC) of activity scores is evaluated for each pair of regulons. Next, for a fixed pair of regulons, A and B, the corresponding CSI is defined as the fraction of regulons whose PCC with A and B is lower than the PCC between A and B.

Hierarchical clustering with Euclidean distance was performed based on CSI matrix to identify different regulon modules. We also use CSI>0.7 as cutoff to build the regulon association network to investigate the relationship of different regulons and visualized it by Cytoscape (Shannon et al., 2003). We used the same strategy to identify submodules within M7. For each regulon module, its activity score associated with a cell type is defined as the average of the activity scores of its regulon members in all cells within this cell type. Then the top ranked cell types are identified for each module.

### Quantifying cell type relationship

Using the gene regulatory network analysis as a guide, we quantified the relationship between different cell-types based on the similarity of the overall regulon activities, which is quantified by the Spearman correlation coefficient. The results were visualized as a network, where a pair of cell types were connected if the Spearman correlation coefficient is greater than 0.8. Groups of related cell-types were identified by using the Markov Clustering Algorithm (MCL) (van Dongen and Abreu-Goodger, 2012), as implemented in the ClusterMaker application in Cytoscape. We used the default setting except setting the inflation parameter as 2.

### Web service

We created an interactive, web-based portal to explore the network atlas in this study (URL: http://regulon.rc.fas.harvard.edu). This interactive website is constructed with some of latest technologies including JavaScript libraries jQuery 3.3, Bootstrap 4, and Leaflet 1.3. Together these libraries provide efficient client-side search, zooming functions for the large cell type network. The site is hosted on an Apache web server running the Apache Tomcat which provides the necessary back-end support for the web server. Users can zoom-in on a part of network, mouse-over, click on a cell type in the network, and browse information about the associated regulons and other most similar cell types. The website also provides a complete, downloadable list of pairwise regulon-cell type associations.

## Author contributions

G.C.Y. conceived the study. S.S. performed computational analysis. Q.Z. built the web service.S. analyzed cell-type network. L.F. helped with raw data processing. G.G. discussed the manuscript and provided input about the MCA data. S.S., G.C.Y., Q.Z. and A.S. wrote the manuscript. All authors contributed ideas for this work. This research was supported by a Claudia Barr Award and NIH grant R01HL119099 to G.C.Y.

## Competing interests

The authors declare no competing financial interests.

## Figure legends

**Figure S1**. Comparison of our Avg20 approach with the original SCENIC implementation. (A) Bladder; (B) Kidney; (C) Bone marrow. For each tissue, the comparison is done in several ways. Left panels: comparison of RAS-based silhouette values. The Avg20 approach was repeated three times. Top right panels: overlap of detected regulons between different Avg20 replicates. Bottom right panels: correlation between different Avg20 replicates based on pairwise distance of all single cells in each tissue.

**Figure S2**. Evaluating the effect of grouping size on the performance of SCENIC analysis. The analysis is based on sampled single cells (∼61k) from multiple tissues. (A) Overlap of regulons identified based on different Avg20 replicates and their correlations. (B) Overlap of regulons identified based on pooled data with different group sizes. (C) Comparison of silhouette values which calculated based on regulon activity matrixes derived from Avg20, Avg50 and Avg100 profiles.

**Figure S3**. t-SNE map for all sampled single cells (∼61k) based on RAS. Each cell is color-coded based on tissue (A, B) or cell type assignment (C). (A) All sampled single cells are highlighted. (B) Cells obtained from each individual tissue are highlighted. (C) Cells obtained from each specific cell type are highlighted.

**Figure S4**. Cell-type specific regulon activity analysis. Similar to Figure 2 but for additional cell types.

**Figure S5**. Activation of regulon modules in different cell types. (A) Regulon association network based on CSI matrix (CSI>0.7). Different colors represent different regulon modules.(B) Average activity scores of 8 representative regulon modules in different cell types. (C) Average module activity scores mapped on t-SNE.

**Figure S6**. Relationship network for the 98 major cell types. Each node represents a group of 20 cells within the same cell-type and tissue.

## Supplementary table legend

**Table S1**. The TFs and their target genes for all 202 regulons, each row means one regluon.

**Table S2**. Cluster numbers and the corresponding definition of cell type identities for 98 major cell types.

**Table S3**. Average activity scores for all regulons in 98 major cell types, each row means each regulon and each column means each major cell type.

**Table S4**. Specificity scores for all regulons in 98 major cell types, each row means each regulon and each column means each major cell type.

**Table S5**. Cell type related GEO datasets in which the target genes of regulons are significantly co-expressed based on SEEK analysis.

**Table S6**. 8 regulon modules and detailed regulon names contained in each module.

**Table S7**. Cell type cluster number of each tissue, number of single cells and the corresponding definition of cell type identities for 818 cell types.

**Table S8**. Average activity scores for all regulons in 818 cell types identified from the whole MCA dataset, each row means each regulon and each column means each cell type.

**Table S9**. Specificity scores for all regulons in 818 cell types identified from the whole MCA dataset, each row means each regulon and each column means each cell type.

